# Temporal anticipation based on memory

**DOI:** 10.1101/090555

**Authors:** André M. Cravo, Gustavo Rohenkohl, Karin Moreira Santos, Anna C. Nobre

**Affiliations:** Centro de Matemática, Computação Congnição, Universidade Federal do ABC (UFABC); Department of Experimental Psychology and Oxford Centre for Human Brain Activity, University of Oxford

## Abstract

The fundamental role that our long-term memories play in guiding perception is increasingly recognised, but the functional and neural mechanisms are just beginning to be explored. Though experimental approaches are being developed to investigate the influence of long-term memories on perception, these remain mostly static and neglect their temporal and dynamic nature. Here we show we show that our long-term memories can guide attention proactively and dynamically based on learned temporal associations. Across two experiments we found that detection and discrimination of targets appearing within previously learned contexts are enhanced when the timing of target appearance matches the learned temporal contingency. Neural markers of temporal preparation revealed that the learned temporal associations trigger specific temporal predictions. Our findings emphasize the ecological role that memories play in predicting and preparing perception of anticipated events, calling for revision of the usual conceptualisation of contextual associative memory as a reflective and retroactive function.

---

Perception is increasingly recognised to be a highly proactive process resulting in a selective (re)construction of the external milieu that emphasises items and attributes that may be adaptive in a given context. Goal-driven selective attention has provided a successful paradigm for investigating the sources and mechanisms of top-down modulation of signal processing within perceptual streams. Decades of research have yielded enormous progress in revealing how the locations and feature-related attributes of relevant events are prioritised and integrated along the sensory hierarchies (Desimone & Ducan 1995; Fries 2015; Kastner & Ungerleider 2000; Reynolds & Chelazzi 2004). These top-down biases were subsequently shown also to carry dynamic information about the estimated timing of relevant events – a phenomenon called temporal orienting of attention or, more generally, temporal expectation (Nobre & Rohenkohl 2014). Trying to understand how temporal predictions of relevant events are extracted and can guide top-down control has become an active area of research, with promising inroads being made (Calderone et al. 2014; Cravo et al. 2013; Doherty et al. 2005; Lakatos et al. 2008; Rohenkohl & Nobre 2011; Vangkilde et al. 2005).

As the attention field matures, scholars have returned to older hypothesised sources of top-down control of perception. In addition to current goals uploaded into short-term stores, our long-term memories have been proposed to guide perception from the earliest days of empirical psychology (Helmholtz 1867). Contemporary research using various types of tasks vindicates this classic notion(Goldfarb et al. 2016; Chun 2000; Giesbrecht et al. 2013; Hutchinson & Turk-Browne 2012; Kasper et al. 2015; Kunar et al. 2008; Stokes et al. 2012; Summerfield et al. 2006; Zhao et al. 2013). The tasks used, however, tend to focus on static aspects of learned contingencies, such as the target location or identity. In the current study, we asked whether our long-term memories can also carry temporal information that can guide perceptual analysis proactively and dynamically to enhance the processing of anticipated target attributes at the right time. The research builds on recent discoveries of mechanisms for encoding sequential and temporal information within memory systems (Dragoi & Buzsaki 2006; Eichenbaum 2013; MacDonald et al. 2011; Eradath et al. 2015).

We designed a novel memory-based temporal orienting task, based on previous work in the spatial domain (Stokes et al. 2012; Summerfield et al. 2006; Summerfield et al. 2011) to test for performance benefits conferred by learned temporal associations between target items and complex contexts. In the spatial memory orienting tasks, participants learned unique locations of target items embedded within complex scenes during a *learning* session. In a subsequent memory-guided *orienting* session, the scenes served as memory cues for the location of target appearance. Perceptual sensitivity and reaction times were strongly facilitated for targets appearing at the learned location relative to an unlearned location (Stokes et al. 2012; Summerfield et al. 2006; Summerfield et al. 2011)

In the current study, participants learn that the target event occurs after a specific temporal interval within a given context - Short (800 ms) or Long (2000 ms). They subsequently perform a memory-based temporal orienting task in which they are asked to detect (Experiment 1) or discriminate (Experiment 2) the target appearance in the studied contexts.

During the Learning task (Figure 1) of both experiments, participants viewed 96 scenes repeated in random order over five (Experiment 1) or seven (Experiment 2) blocks and learned the temporal interval at which the target event occurred within each scene. In each scene, a placeholder black bomb appeared after 1500 ms. The target event was the brief ‘activation’ of the bomb stimulus after either 800 or 2000 ms. In Experiment 1, participants made a simple detection response if the bomb turned blue (80% of trials), but withheld responding if the bomb turned red (20%). In Experiment 2, participants made a forced-choice response depending on whether the bomb turned blue or green (50% each). Participants had to respond to the target event within a short RT window (600 ms) in order to prevent the bomb from exploding. If participants responded correctly a smoky cloud was presented, indicating that the response was correct. If participants did not respond, an explosion image was presented.

We measured the benefits of learning the temporal relationship between scenes and target by comparing reaction times to the targets from the first and last block of the Learning session. In Experiment 1, participants had better performance at the end of the Learning session for both Short and Long intervals (two-way Interval × Block ANOVA: main effect of Interval, F_1,9_=50.74, P<0.001, η^2^_*partial*_=0.435, main effect of Block, F_1, 9_=105.79, P<0.001, η^2^_*partial*_=0.852, interaction, F_1,9_=7.56, P=0.02, η^2^_*partial*_=0.063). However, learning was stronger for scenes with Short intervals (t_9_=2.75, P=0.02, *d*=0.869). For Experiment 2, benefits in performance depended on the interval (two-way Interval × Block ANOVA: main effect of Interval, F_1,13_=9.25, P=0.009, η^2^_*partial*_=0.016, no main effect of Block, F_1,13_=3.04, P=0.11, η^2^_*partial*_=0.063, interaction, F_1,13_=5.09, P=0.04, η^2^_*partial*_=0.007). Specifically, reaction times improved only for Short intervals (first versus last block for Short intervals, t_13_=2.85, P=0.014, *d*=0.762, Long intervals, t_13_=0.74, P=0.47, *d*=0.198). Thus, in both experiments, systematic decreases in reaction times suggested that participants learned the temporal relationship between scenes and target intervals, with more pronounced learning for the short interval, as expected according to the hazard effect (Nobre & Rohenkohl 2014; Cravo et al. 2011).

**Figure 1.**
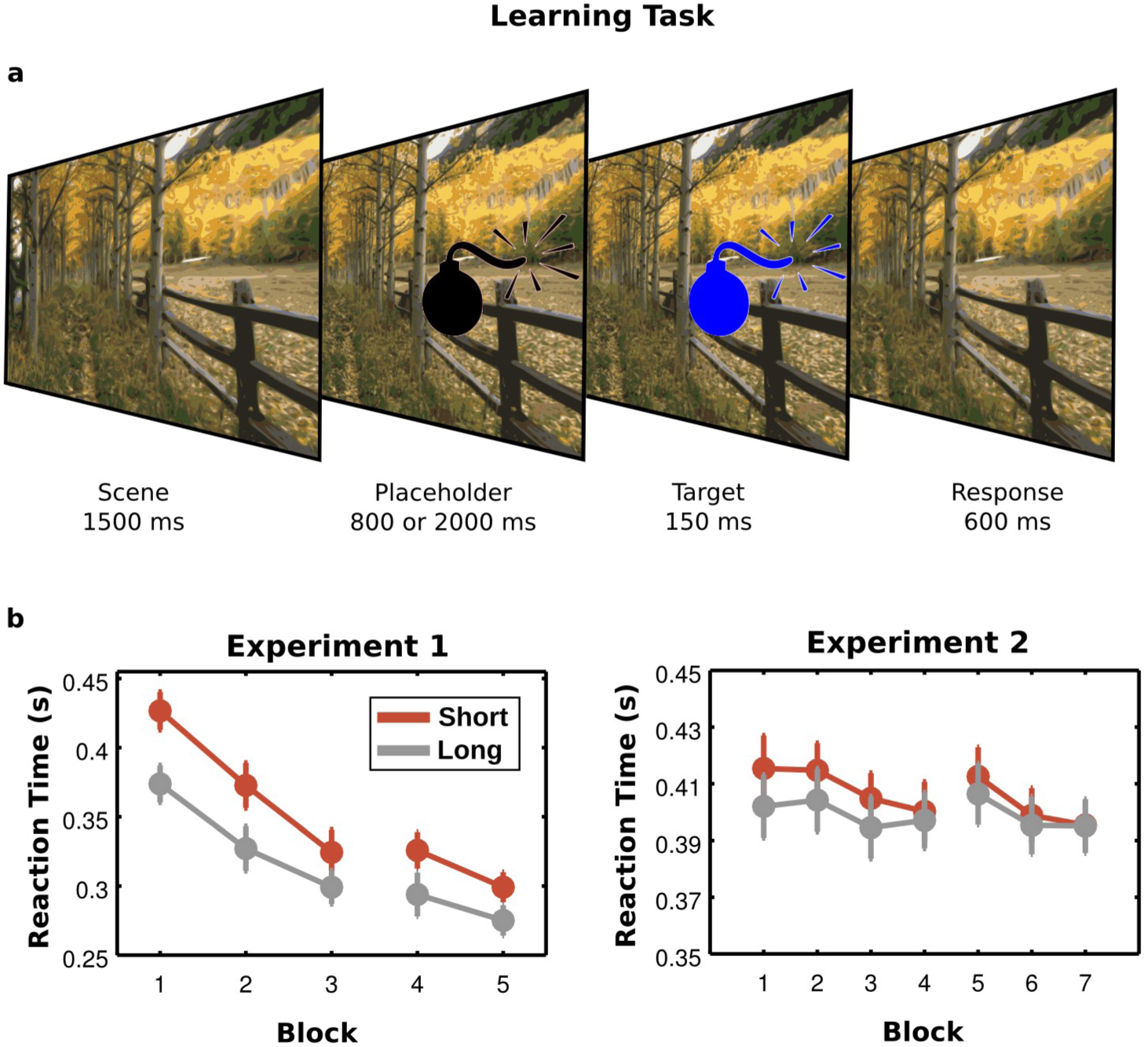
Learning Task. **(a)** During the Learning task, participants viewed a complex scene and learned the temporal interval at which the target event occurred within that scene. After 1500 ms of the scene presentation, a placeholder (bomb) appeared. After an 800-ms (Short) or 2000-ms (Long) interval, the placeholder changed colour. In Experiment 1, the target changed to blue in 80% of trials (Go-target) or red (20% of trials, no-go target). In Experiment 2 the target changed to blue or green in an equal proportion of trials. Participants had to detect the target (Experiment 1) or discriminate the colour of the target (Experiment 2). **(b)** In both tasks, participants’ reaction times decreased as a function of Block, with a stronger effect for Short intervals. All plots show mean and standard error of the mean (SEM) across participants.

The Memory task assessed whether participants formed an explicit memory for the temporal association within each scene (Figure 2). The Memory task was repeated midway through the Learning task (after block 3 in Experiment 1 and after block 4 in Experiment 2) and after completion of the Learning task. During the Memory task, participants viewed each scene in isolation and indicated whether it was associated with a Short or Long interval. In both Experiments there was an increase in accuracy as a function of Learning (two-way Interval × Block ANOVA, Experiment 1: main effect of Block, F_1,8_=20.37, P=0.002, η^2^_*partial*_=0.730, no main effect of Interval, F_1,8_=0.04, P=0.84, η^2^_*partial*_=0.001, no interaction, F_1,8_=0.002, P=0.97, η^2^_*partial*_=0; Experiment 2: main effect of Block, F_1,13_=23.02, P<0.001, η^2^_*partial*_=0.352, no main effect of Interval, F_1,13_=3.74, P=0.075, η^2^_*partial*_=0.065, no interaction, F_1,13_=0.269, P=0.613, η^2^_*partial*_=0.001). The results showed that participants formed reliable explicit memories for the temporal associations between scenes and target presentation (Figure 2).

**Figure 2.**
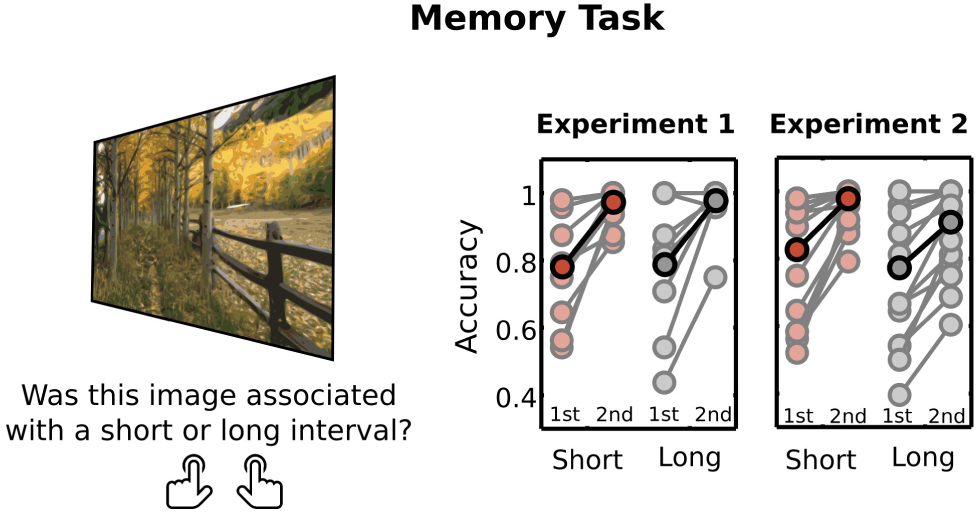
Memory Task. In the Memory task, participants viewed each scene in isolation and indicated whether it was associated with a Short or Long interval. This task was performed by the participants halfway through the experimental session (1^st^ session) and at the end of the Learning task (2^nd^ session). Mean accuracies show how participants improved their performance over Learning blocks, forming reliable explicit memories for the temporal associations between scenes and target presentation. All plots show mean accuracy across participants (darker colours) and raw data from all participants (lighter colours)

The final Orienting task probed whether the learned temporal associations influenced behavioural performance to expected targets. Whereas standard temporal orienting tasks (Miniussi et al. 1999; Nobre & Rohenkohl 2014) use predictive symbolic cues, in this memory-guided version only the previously learned temporal associations stored in LTM were available to guide temporal expectation of the target appearance (Figure 3). In the majority of trials, the target occurred at the remembered interval (Valid cue) while in the remaining trials target occurred at the other interval, and the scene thus provided invalid temporal information (Invalid cue).

**Figure 3.**
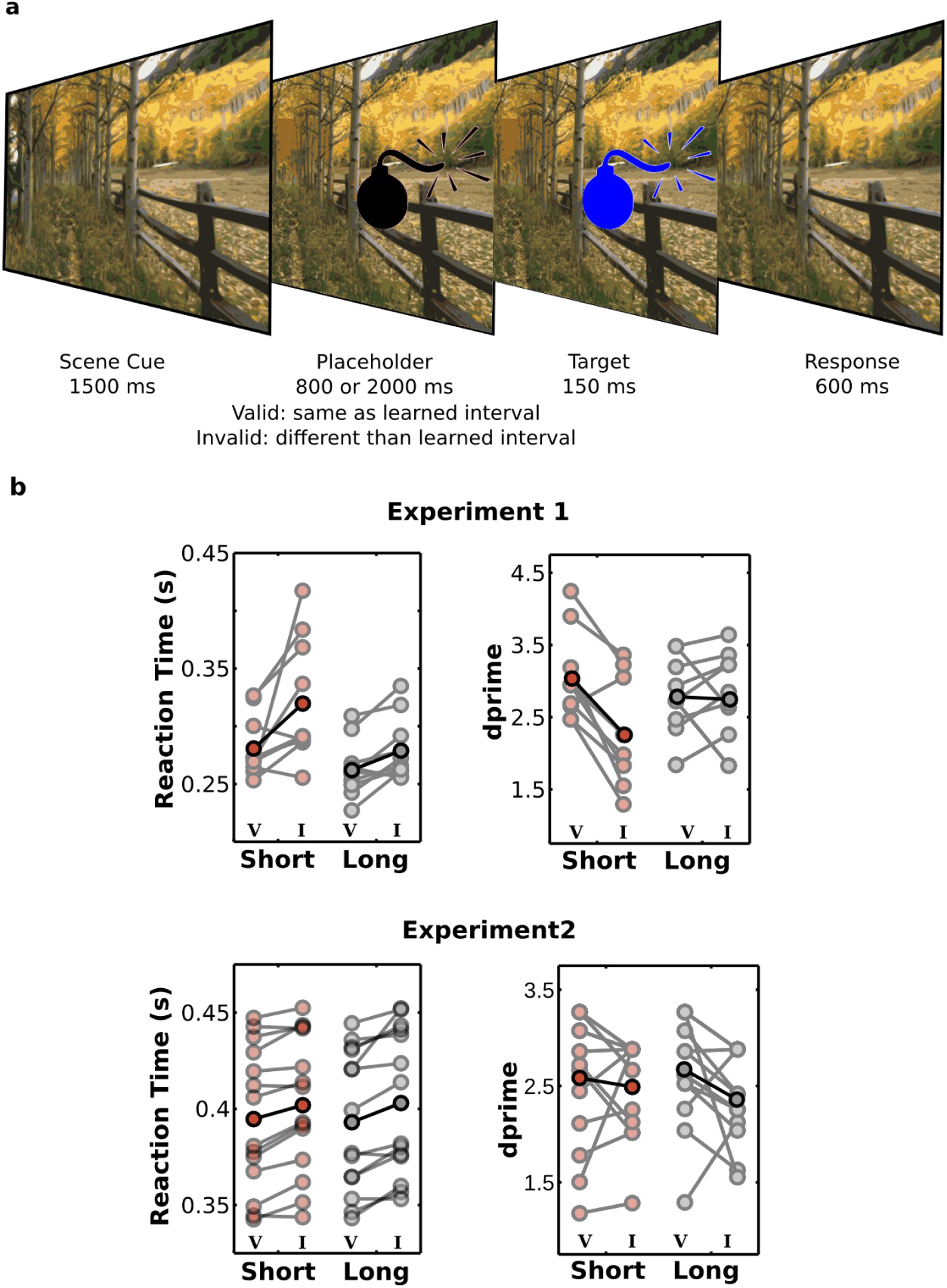
Temporal Orienting Task. **(a)** In the Temporal Orienting task, trial sequence was similar to the Learning task, however the interval when the target appeared matched that in the Learning task in the majority of trials (67%, Valid cues) while in the remaining trials (33%, Invalid cue), the target occurred at the other interval. **(b)** Performance was strongly influenced by long-term memory cues, and both reaction times and perceptual sensitivity were better for Valid (V) than for Invalid (I) scenes. All plots show mean across participants (darker colours) and raw data from all participants (lighter colours).

As shown in Figure 3, performance was strongly influenced by LTM cues. In both experiments, reaction times were shorter when targets were presented at the learned temporal interval (two-way Cue × Interval ANOVA, Experiment 1: main effect of Cue, F_1,9_= 30.47, P<0.001, η^2^_*partial*_=0.290, main effect of Interval, F_1,9_=10.14, P=0.01, η^2^_*partial*_=0.254, no interaction, F_1,9_=2.3, P=0.163, η^2^_*partial*_=0.020; Experiment 2: main effect of Cue, F_1,13_=20.14, P=0.001, η^2^_*partial*_=0.029, no main effect of Interval, F_1,13_=0.42, P=0.530, η^2^_*partial*_=0, no interaction, F_1,13_=0.023, P=0.883, η^2^_*partial*_=0).

Long-term memory also improved perceptual sensitivity, as indexed by d-primes, for both detection (Experiment 1 two-way Cue × Interval ANOVA, main effect of Cue, F_1,9_=9.54, P=0.013, η^2^_*partial*_=0.198, no main effect of Interval, F_1,9_=0.54, P=0.481, η^2^_*partial*_=0.017, interaction, F_1,9_=9.72, P=0.012, η^2^_*partial*_=0.081) and discrimination task (Experiment 2 two-way Cue × Interval ANOVA, main effect of Cue, F_1,13_=7.33, P=0.018, η^2^_*partial*_=0.066, no main effect of Interval, F_1,13_=0.05, P=0.824, η^2^_*partial*_=0.001, no interaction, F_1,13_=0.70, P=0.419, η^2^_*partial*_=0.010). For the detection task, perceptual sensitivity effects were restricted to the Short interval (paired t-test between Valid and Invalid cues for Short intervals, t_9_=4.64, P=0.001, *d*=1.467, for Long intervals, t_9_=0.20, P=0.845, *d*=0.063).

In Experiment 2, EEG data were recorded during the Orienting task and the contingent negative variation (CNV) provided a neural marker of proactive temporal expectation (Cravo et al. 2011; Miniussi et al. 1999; Praamstra et al. 2006). The CNV for Valid cues had higher (more negative) amplitudes for the period from 90 ms to 340 ms after cue presentation (cluster-stat=202.05, cluster-P=0.002) and for the period 390 ms to 800 ms after cue presentation (cluster-stat=363.30, cluster P<0.001).

The steeper development of the CNV for scenes associated with short than with long intervals (Figure 4) paralleled how temporal expectations influence neural preparation in perceptual tasks using symbolic cues (Cravo et al. 2011; Los & Heslenfeld 2005; Miniussi et al. 1999). Importantly, the amplitude of the CNV correlated significantly with response times, indicating a functional relation between neural preparation and behavioural performance (t_13_=2.69, P=0.018, *d*=0.719 Figure 4c).

An important property of learned temporal contextual associations is that their strength can vary. Participants formed stronger temporal memories for some scenes than for others as shown by the association between response time and accuracy during the memory test (t-test on the estimated slopes, t_13_=-3.53, P=0.004, *d*=0.943 Figure 4d). We took advantage of this variability to investigate whether the quality of temporal memories influenced the degree of neural preparation and performance benefit. Response times during the Memory task were used as a probe to the strength of the temporal-association memory. As can be seen in Figure 4, Memory Strength was predictive of the CNV amplitude (t-test on the estimated slopes, t_13_=-2.33, P=0.037, *d*=0.620) as well as of behavioural performance benefits (t-test on the estimated slopes, t_13_=-2.71, P=0.018, *d*=0.723).

**Figure 4.**
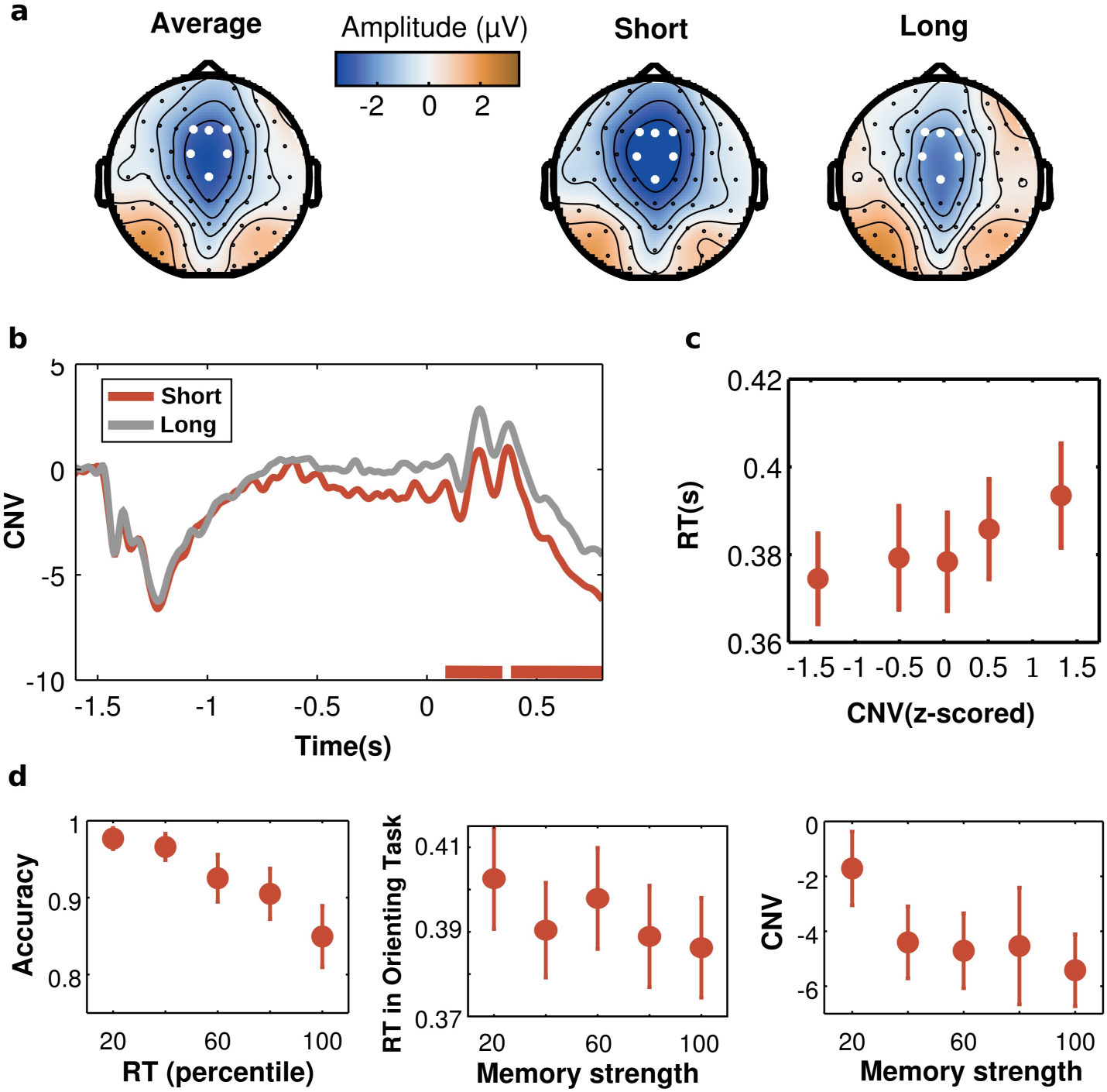
Electrophysiological results. **(a)** Topographies of the grand-averaged Contingent Negative Variation (CNV) and for the CNVs at the Short foreperiod (FP) for scenes associated with Short or Long intervals. **(b)** The CNV recorded during the Orienting Task of Experiment 2 was strongly influenced by the temporal association in memory (red lines at the bottom represent the two temporal clusters where the CNV was larger for Short than for Long temporal associations). **(c)** Larger CNV amplitudes were associated with shorter reaction times. **(d)** Left panel: during the Memory task, shorter response times were associated with higher accuracy. Given this relation, response times were used to create a Memory-Strength index, which estimated the quality of the temporal-association memory. Further analyses showed that stronger memories were associated with shorter reaction times during the subsequent Temporal Orienting task (middle panel) and with CNV amplitude (right panel). All plots show mean and standard error of the mean (SEM) across participants.

Taken together, our results show that long-term memories can guide our perception and behaviour dynamically, utilizing learned temporal associations stored in such memories to prepare neural activity for relevant upcoming events. They cast long-term memories in a new light. Rather than emphasizing their reflective and retroactive role of reconstituting, or re-membering past events; they highlight the proactive role they play in predicting and preparing perception by pre-membering anticipated events. The findings open new lines of investigation into the mechanisms through which mnemonic temporal associations guide perception. A fuller understanding of human perception will require understanding of dynamic regulation by both top-down signals from long-term memories and short-term biases related to current goals and expectations.

## MATERIALS AND METHODS

### Participants

A total of ten volunteers (three female, seven male, mean-age: 19.4) participated in Experiment 1 (Detection) and eighteen (seven female, eleven male, mean-age: 20.17) participated in Experiment 2 (Discrimination). They all gave informed consent. All had normal or corrected vision and were free from psychological or neurological diseases according to self-report. The number of participants was based on comparable sample sizes in the literature (Stokes et al. 2012; Summerfield et al. 2006). The experimental protocol was approved by The Research Ethics Committee of the Federal University of ABC and from the Central University Research Ethics Committee of the University of Oxford.

### Apparatus

The stimuli were created on MATLAB v.7.10 (The MathWorks) and presented using the Psychtoolbox v.3.0 package for MATLAB (Brainard, 1997). Images were displayed on a 21-inch CRT with a spatial resolution of 1024 by 768 pixels and a vertical refresh rate of 60 Hz, placed 100 cm in front of the participant. Responses were collected via a response box (DirectIN High SpeedButton / Empirisoft).

### Stimuli and Task

We conducted two similar experiments, in which participants learned new associations about the timing of a target event occurring within a scene, and then performed an orienting task requiring detection (Experiment 1) or discrimination (Experiment 2) of the target event occurring within the learned context. In Experiment 2, electroencephalography (EEG) activity was recorded during the performance of the final, temporal orienting task requiring target discrimination. Each experiment consisted of three different tasks that took take place on the same day: a Learning Task, a Memory Task, and a Temporal Orienting Task. Participants performed a session of the Learning Task, followed by a Memory Task. They then performed another session of the Learning Task and one more session of the Memory Task. Finally, they performed the Temporal Orienting Task.

### Experiment 1. Detection

#### Learning task

During the learning task, participants viewed 96 complex scenes repeated in random order over five blocks and learned the time for a target event to occur within each scene. Scene stimuli were similar to those used by previous studies (Stokes et al. 2012; Summerfield et al. 2006; Summerfield et al. 2011), consisting of photographs of different indoor or outdoor views. Scenes were prepared using Matlab, and subtended 22° × 17° of visual angle at a viewing distance of 100 cm. Though we considered using dynamic scenes, this would have conflated the timing of the target event with a sequence of spatial and/or feature-related changes that need not specifically rely on learning temporal intervals.

Each scene was associated with a target event being presented in a specific time and place that remained fixed throughout the whole learning session. The target event occurred between 5° to 7° of visual angle along both the lateral and longitudinal axes and was preceded by a placeholder presented at the exact same location. Participants were instructed to learn when the target event was presented within each scene. The interval and location of the target within each scene was randomised between participants.

Each trial started with the presentation of one of the scenes and a Fixation Cue in the centre of the screen. After a period of 1.5 seconds, a placeholder black bomb (1° x 1°) was presented in either the upper or lower quadrant of the right or left side of the scene. After an interval of either 800 ms or 2000 ms, the bomb changed its colour to blue (Go target, 80% of the trials) or red (No-Go target, 20% of the trials). The type of target (Go or No-Go) was randomised over scenes and participants were instructed that the same scene could have Go or No-Go targets in different blocks. Half of the images (48 scenes) were associated with each interval (Short or Long). Participants were instructed to respond as quickly as possible to Go targets. If participants responded correctly and quicker than 600 ms, a smoky cloud was presented, indicating that the response was correct. If participants did not respond to Go targets within 600 ms or if they responded to No-Go targets an explosion image was presented. The order of scene presentation was randomized in each block. Participants performed three learning blocks in a row and then performed a Memory task. They then completed two more learning blocks followed by another Memory Task.

#### Memory Task

During the memory task, participants viewed the same 96 naturalistic scenes repeated in random order. The scenes were presented on their own (no bombs appeared), and remained on the screen until participants responded. Their task was to indicate if the scene was associated with a Short (800 ms) or Long (2000 ms) interval during the learning task. Responses were made using index/middle fingers of the right hand. Memory tasks were performed after three blocks of the Learning task and after the final block of the Learning task.

#### Temporal Orienting Task

After completing five blocks of the Learning task and two Memory tasks, participants performed the Temporal Orienting Task. The task was similar in structure to the Learning Task. Participants viewed the same 96 scenes, in which a bomb changed colour after a Short or Long interval. In the majority of the trials (67%), the interval in the Orienting task was the same as the learned interval in the Learning Task. The scene therefore triggered a valid memory cue for target timing. In the remaining trials (33%), the interval was switched, and the scene provided an invalid temporal memory cue. As before, participants were instructed to respond as quickly as possible to Go targets and to withhold responding to No-Go targets. The Temporal Orienting task consisted of three blocks, each with 96 scenes. In each block, a different subset of the scenes were selected to have an invalid memory cue. No feedback (smoky cloud or explosion) was given during this task.

### Experiment 2. Discrimination

The second experiment served as a replication and extension of Experiment 1, with EEG recordings made during the Orienting Task. The experiment contained the same three phases. The major differences were that instead of using Go/No-Go targets, a change in bomb colour (blue or green) required a discrimination response. Participants were instructed to press the right button when the bomb turned blue and the left button when it turned green (the mapping of colour and response was counterbalanced across participants). Blue and green bombs were equiprobable and occurred arbitrarily for each scene. Participants were instructed that each scene was associated with the target event being presented in a specific time and place, but that there was no association between the scene and the colour of the bomb. Instead of performing five Learning blocks as in Experiment 1, participants performed seven Learning blocks. The Memory task was performed after four blocks of Learning and then after the final Learning Task block. The Temporal Orienting task was performed last.

### EEG recording and pre-processing

Continuous recording from 64 ActiCap electrodes (Brain Products, München, Germany) at 1000 Hz referenced to FCz (AFz ground) provided the EEG signal. The electrodes were positioned according to the International 10–10 system. Additional bipolar electrodes recorded the EOG. EOG electrodes were placed to the side of each eye (HEOG) and above and below the right eye (VEOG). EEG was recorded using a QuickAmp amplifier and preprocessed using BrainVision Analyzer (Brain Products). Data were downsampled to 250 Hz and re-referenced to the averaged earlobes. To remove eye-blink artifacts, filtered data (0.05–30 Hz) were subjected to independent component analysis. Eye-related components were identified through comparison of individual components with EOG channels and through visual inspection. Vertical eye activity was removed using ICA.

For the CNV analyses, epochs were created segmented from 250 ms before scene onset until 800 ms after cue presentation. Epochs containing excessive noise or drift (±100 μV at any electrode) or eye artefacts (saccades) were rejected. Saccades were identified as large deflections (±50 μV) in the horizontal EOG electrodes. All data were subsequently checked by visual inspection. Data from four participants were removed due to excessive eye-movements (two participants) or an excessive number of rejected trials (two participants). A small proportion of trials of the remaining participants were rejected (0.05 ± 0.01). We focused our analyses on Short-Valid and Long-Valid cues, with an average of around 90 clean epochs per condition.

### Behavioural Analyses

To quantify the improvement in performance in the Learning tasks, RTs from the first and last blocks for Short and Long intervals were submitted to a 2x2 repeated-measures ANOVA, with factors Interval (Short x Long) and Block (First x Last).

For the Memory task, mean accuracy for scenes with Short and Long intervals for the two blocks of the Memory Task were submitted to a repeated-measures ANOVA, with factors Interval (Short x Long) and Block (First x Last). The data from one participant for the second Memory session of Experiment 1 was not saved due to technical problems. To perform the repeated-measures ANOVA, all data from this participant was removed for this particular analysis.

For the Temporal Orienting Task, mean RTs for correct responses were submitted to a repeated-measures ANOVA with Interval (Short x Long) and Cue (Valid x Invalid) as factors. We further calculated d-primes for each condition in the Temporal Orienting Task. In Experiment 1, hits were considered as a correct response for a Go target while false alarms were considered when participants responded to a No-Go target. D-primes were submitted to a repeated-measures ANOVA with Interval (Short x Long) and Cue (Valid x Invalid) as factors. In Experiment 2, hits were calculated as correct response for green targets and false alarms as incorrect responses for blue targets. D-primes were submitted to a repeated-measures ANOVA with Interval (Short x Long) and Cue (Valid x Invalid) as factors.

### Contingent Negative Variation (CNV)

In the Orienting task of Experiment 2, analyses of the CNV focused in central-midline electrodes (F1/Fz/F2/FC1/FC2) for scenes associated with a Short and Long intervals during the Learning Task. A cluster-based analysis (Maris & Oostenveld 2007) was used to compare the CNV between conditions for the time period between scene presentation and the first possible moment of the target. The nonparametric statistics were performed by calculating a permutation test in which experimental conditions were randomly intermixed within each participant and repeated 1000 times.

To test whether the CNV reflected a stronger temporal anticipation, we investigated if there was a relation between CNV at the single-trial level and RTs. This analysis was performed in scenes associated with short intervals in the Learning task and that were presented at the short interval in the Temporal Orienting task (Short Valid cues). The CNV activity for the second cluster (from 390 ms to 800 ms after cue onset) was averaged for each trial, z-scored and separated into five bins (each with 20% of the data). The associated RT for each bin was calculated and a nonparametric regression was calculated for each participant. At the group level, the Fisher transformed estimated coefficients for the regression were compared to zero using a t-test.

### Memory-strength index

To estimate the strength of the temporal-association memories, we used the response times during the Memory task. In a first step we investigated whether these response times were correlated with response accuracy. For each participant, response times for all scenes during the second Memory task (after completion of the Learning task) were separated into 5 bins, each containing 20% of the data. Response times shorter or longer than 2.5 standard deviations were removed prior to binning. For each bin, the mean accuracy was calculated. A nonparametric regression was performed separately for each participant. At the group level, the Fisher transformed estimated coefficients were compared to zero using a paired t-test.

Given the strong association between response time and accuracy, we used these responses times as a Memory Strength index in two following analyses. In a first analysis, we investigated whether this index was associated with shorter reaction times in the subsequent Temporal Orienting task. If participants had a stronger association between a given scene and its learned interval, then they should benefit more strongly from this association. We focused our analysis on: (1) the first block of the Temporal Orienting task; (2) Short Valid trials; (3) Trials in which participants gave correct responses in the Temporal Orienting task; (4) Scenes that participants judged correctly in the Memory task. These restrictions were used to isolate as maximally as possible the effect of Memory on performance.

For each trial in the Temporal Orienting task conforming to the above-mentioned restrictions, the response time for that scene in the Memory task was used as a predictor of the RT in the Temporal Orienting task. The Memory strength index was calculated as the percentage of response times that were longer than each individual response time. For example, for the shortest response time, all other response times were longer, resulting in a Memory Strength index of 100. A nonparametric regression was performed with the RT in the Temporal Orienting task as the dependent variable and with the Memory Strength index as the predictor. At the group level, the Fisher transformed estimated coefficients were compared to zero using a paired t-test.

A similar analysis was performed to test whether this index was also related to the CNV. The same restrictions were used and the Memory Strength index was calculated in a similar way. The CNV was measured in the same electrodes as previously mentioned and in the time period of the second significant cluster (390 ms to 800 ms). A nonparametric regression was performed with the CNV as the dependent variable and with the Memory Strength index as the predictor. At the group level, the Fisher transformed estimated coefficients were compared to zero using a paired t-test.

